# A framework to assess the effects of changes in species composition on processes derived from trophic interactions

**DOI:** 10.1101/2020.11.09.374389

**Authors:** Anna R. Landim, Fernando A. S. Fernandez, André T. C. Dias

## Abstract

Functional diversity uses response and effect traits to understand how communities are affected by changes in the environment and their consequences on the structure and functioning of ecosystems. However, most studies focus on a single taxonomic or functional group, ignoring that many ecological processes result from trophic interactions. Here we established a multi-trophic trait-based framework to evaluate the consequences of community change for ecological processes resulting from trophic interactions. Specifically, we estimated the potential effect of each species considering the consumer and resource communities involved on the trophic interaction. The functional space of consumer and resource communities were incorporated into a single analysis by using resource traits to estimate consumers’ functional space. Our framework included a parameter that establishes different weights to unique interactions when estimating a species potential effect. We presented two modifications for application using abundance and species richness data and two modifications to allow incorporating absent species into the analysis. Our framework can be used to investigate consequences of community changes in different situations, such as species extinctions, invasions and refaunation. To demonstrate the insights derived from our framework we used an exemplary study case of refaunation of an impacted tropical forest. Our framework informs on a species contribution to an ecological process according to its originality, i.e., the uniqueness or redundancy of its interactions, and the magnitude of the effect, indicated by the frequency of the resource’s community trait values with which it interacts. Thus, it helps to increase the understanding of the effects of changes in community composition on ecological processes resulting from trophic interactions. It assists practitioners and researches with predictions and evaluations on potential loss and reestablishment of ecological functions resulted from changes in community functional composition.

## 1 Introduction

Community composition is dynamic (Connell and Sousa 1983), always changing over evolutionary and ecological time. In the Anthropocene, human activities have intensified and accelerated these changes; species are lost and added through extinction and colonization with potential consequences to ecosystem functioning and services delivery (Bommarco et al. 2013, Malhi et al. 2016). Changes in species composition occur both locally and regionally, and modified communities often do not provide the same ecosystem functions as compared to the original ones (Dornelas et al. 2014). Many ecosystem processes are determined by the combined effect of traits from species belonging to different trophic groups (de Bello et al. 2010). Ecological processes resulting from trophic interactions are particularly vulnerable because species from different trophic levels often show different responses to environmental changes (Berg et al. 2010). This can result in mismatches between consumer and resource species (Törnross et al. 2018, Nowak et al. 2019), with detrimental consequences to ecosystem functions.

Functional traits are being increasingly used to asses species niches’ dimensions (Violle and Jiang 2009), as traits can reflect both how species respond to environmental changes (Grinnellian niche) and how species affect ecosystem processes and other species (Eltoninan niche) (Rosado et al. 2016). In this sense, functional traits are used to understand how communities respond to environmental changes (Nowak et al. 2019) and how the consequent shifts in community composition affects ecosystem functioning (Suding et al. 2008, Cadotte et al. 2011). Trait-based perspectives, rather than taxonomic ones, have been encouraged, because traits facilitate detecting patterns and making predictions, while taxonomy do not directly reflect functionality (Díaz et al. 2006). In this sense, communities can strongly differ in species composition but still show very similar functioning (Dehling et al. 2020).

In the last years, several conceptual and analytical frameworks linking response and effect traits have emerged. Initially, most frameworks considered just one taxon (Lavorel and Garnier 2002, Suding et al. 2008, Gross et al. 2009, Wallenstein and Hall 2012). However, many ecosystem processes derive from interactions between species belonging to different trophic groups (e.g. pollination and seed dispersal) (Akçakaya et al. 2019). Additionally, the effect of different trophic levels on each other has influence on ecosystem function (Moretti et al. 2013). Thus, there is a concern on incorporating more than one trophic level into response-effect frameworks (Lavorel et al. 2013, Schleuning et al. 2015). However, functional groups, like pollinators, can be locally composed by a wide range of taxonomical groups as distinct as lizards, birds, wasps, flies, beetles and butterflies (Kaiser-Bunbury et al. 2017). These taxonomical groups may exhibit quite different traits and strategies by which they assess their resources, making it difficult to select relevant traits across taxa and, therefore, include all taxa engaged in the interaction in a single analysis.

When evaluating an ecological process derived from trophic interactions, a consumer’s role can be assessed by a set of trait values from the resource with which it is able to interact (Dehling et al. 2016). The process-related niche, proposed by Dehling and Stouffer (2018), uses resource traits to build consumers’ functional space. This allows the combination of taxonomic groups which differ in morphological traits, but share the same function in an ecological interaction. For instance, in the example of pollinators cited above, small bees can enter and pollinate flowers with long corolla tubes, similarly to large butterflies with long proboscids. Thus, quantifying the traits of the flowers with which each pollinator interacts might result in a better inference of its role in the interaction network. This means that the focus is shifted from the comparison of trait values between species, to the role of different species on processes. Additionally, using the same traits to characterize consumers’ and resources’ functional spaces supports the incorporation of both trophic levels into the same model.

For predicting the effect of a given species in an ecosystem process, it is fundamental to understand its originality, which can be defined as a species contribution to its community functional space. That is, the set of shared (redundant) and unshared (unique) traits with other species in the community (Kondratyeva et al. 2019). The more different is the species’ functional space from the rest of its community, the greater the value of its originality. Additionally, we consider the resource availability, represented by the resource community’s functional space, an important variable to estimate the magnitude of a species potential effect on trophic interactions. Species with the same value of originality will have different potential effects if one interacts with more frequent trait values from the resource community than the other. Thus, we argue that the effect of extinction or addition of a species in an ecological process is defined by its originality and the frequency of traits of the resources with which it interacts.

Here, we use the concept of process-related niche (Dehling and Stouffer 2018) to build an analytical framework to predict the effect of species extinctions and additions on ecological processes resulting from trophic interactions. First, we present an estimate for a species’ originality regarding their community functional space. Then, we propose a measure for the species potential effect on the ecological process being analysed, based on the functional space of the other community involved in the interaction. We also present an estimate for niche amplitude, which helps to interpret the analysis here proposed, as it is one of the variables determining the contribution of a species to its community functional space. Finally, we provide modifications allowing the use of different types of community data, making it easier to use the framework according to data availability.

## 2 Methods

Our model is based on the trait probability density (TPD) framework proposed by Carmona et al. (2016). TPD was created to investigate the relation between traits and fitness and considers the heterogeneous distribution of species fitness across the niche. Another asset is that it allows working with multidimensional trait spaces. Considering the process-related niche (Dehling and Stouffer 2018) the TPD accounts for resource preference, as the consumption probability is dependent on resource traits. Thus, it gives the probability of a species consume different trait values.

We followed Carmona et al. (2016) for building trait probability densities of individuals, species and communities, termed TPD_*i*_, TPD_*s*_ and TPD_*c*_, respectively. Ideally, first the TPD_*i*_ should be estimated. The average across all individuals’ TPD_*i*_ from a given population or species results in the population or species TPD_*s*_. Similarly the weighted average of the co-occurring species TPD_*s*_ result on the community TPD_*c*_. The weights are given by the species’ relative abundances.

Among the options for estimating TPD, the kernel density estimate is the most common one and was used to create this framework using the *ks* package (Duong et al. 2020). Kernel density functions are non-parametric, allowing to estimate irregular trait functions. However, when there is no data to build trait distribution, the TPD can also be estimated through individuals or species mean trait values and their variance (Carmona et al. 2016). The TPD of species and communities were estimated and analysed using R 3.6.2 (R Development Core Team 2019).

## 3 A framework to estimate species effects on ecological processes

Studies that analyze species effects on ecological processes usually consider the functional trait space of only one of the communities (e.g. resources or, consumers) involved in the interaction to determine species roles (Finerty et al. 2016, Dehling et al. 2020). We argue that for a full comprehension of ecological processes involving trophic interactions, both communities must be included. The traits of the resource community represent the availability of potential interactions, while the traits of the resource consumed by the consumers community represent the realization of these interactions (Figure 1).

**Figure 1:**
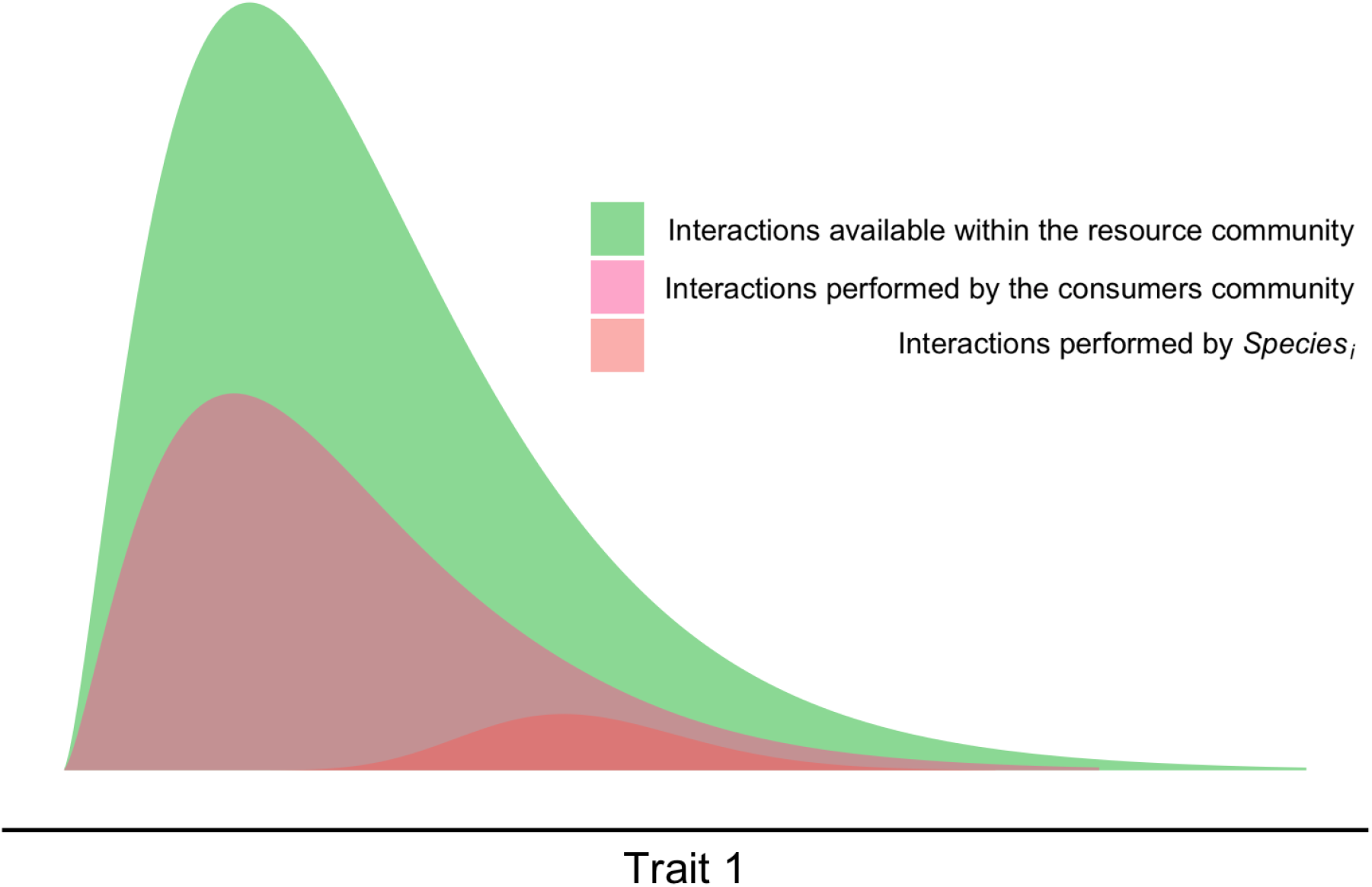
Conceptual model showing the functional space of the communities. In green is depicted the resources community C_ℜ_ that represents the available niche space for potential interactions. In pink, the functional space of the consumers community C_ℭ_ represents the portion of the resource functional space that can be consumed. In red, is represented the functional space occupied by the species i, which is a member of the consumers community.

Species roles on an ecological process is usually estimated by comparing a species functional niche to the rest of its community. The result of this analysis does not allow predictions, as it represents the probability of species participating in an interaction given that this interaction has occurred. To predict the probability of a consumer species participating in an interaction, we need to consider the availability of the resources with which it interacts.

We represent by *Y*_*i*,𝔞_ the effect of species *i* from the community 𝔞 on the ecological process *Y* under investigation. The effect depends on the resource availability represented by the functional space of the resources community, and on the species functional originality (we follow the definition of Kondratyeva et al. (2019) on functional originality). Here 𝔞 stands for the consumers community, represented by ℭ, or the resources community, represented by ℜ.

The functional space of a community, denoted by *C*_𝔞_, is composed by the assembly of the functional spaces of the totality of its species (*N, see Equation 1*). The contribution of a species *i* to its community is given by the product of its 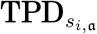 and its relative quantitative contribution to the interactions *V*_*i*,𝔞_*/V*_𝔞_. *V*_𝔞_ represents the total number of interactions in the community 𝔞 and *V*_*i*,𝔞_*/V*_𝔞_ the relative quantitative contribution of the species *i* to the ecological process under scrutiny. *C*_𝔞_ is a TPD_*c*_, which means it is positive and its integral is equal to 1.

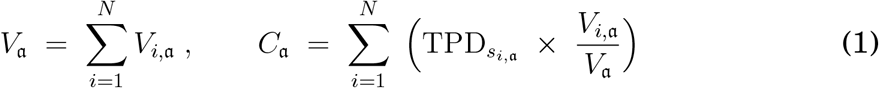

Note that *C*_𝔞_ and 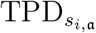 are functions of the traits, even if not explicitly indicated in the notation. If we consider seed dispersal, for example, for a resource species *i, V*_*i*,ℜ_ provides the total amount of seeds produced by species *i* during a given time. This volume multiplied by 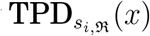 represents the amount of seeds with trait value *x* produced by species *i*. Similarly, for a consumer species *j, V*_*j*,ℭ_ multiplied by 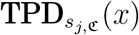 stands for the total amount of seeds with trait value *x* consumed by species *j* in the same period of time.

The species originality, denoted by *O*_*i*,𝔞_, is the full contribution of species *i* to the trait value diversity of its community. It is a result of the balance between the species’ uniqueness and redundancy. Uniqueness expresses the species set of trait values that are not shared with other species in the community, while redundancy is the set of shared trait values (Kondratyeva et al. 2019). The originality of species *i* is estimated by dividing its functional space by the community’s functional space:

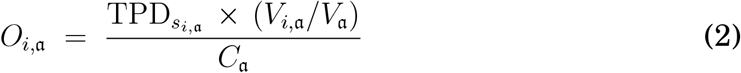

In this sense, *O*_*i*,ℭ_(*x*) represents the proportion of the functional space from the consumers community occupied by species *i* (equation (2)). Note that it is a positive function bounded by 1. The closer it gets to 1, the more unique species *i* is regarding trait value *x*. Species with higher originality have more unique trait values than species with lower originality.

A parameter *θ* ≥ 1 is included in the framework to accentuate species originality in relation to the studied interactions. We define a consumer species effect *Y*_*i*,ℭ_ by the equation 3:

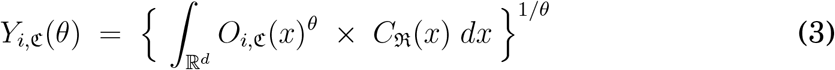

In this equation, *C*_ℜ_(*x*) *dx*, introduced in equation (1), stands for the probability of finding an individual with trait value *x* in the resource community. Given that an individual with trait value *x* was consumed, *O*_*i*,ℭ_(*x*) is the probability that the consumer was an individual from species *i*, ℭ. *C*_ℜ_(*x*) *dx* provides the magnitude of species *i* potential effect. The higher the frequency of trait values that species *i* interacts with, the higher the magnitude of its effect will be. *Y*_*i*,ℭ_(*θ*) indicates the proportion of the functional space from the consumer community occupied by species *i* [to the power *θ*] weighted by the resource availability.

The species *i* potential effect, *Y*_*i*,ℭ_(*θ*), varies between 0 and 1 and is an increasing function of the parameter *θ*. When *θ* increases to infinity, the potential effect converges to the maximum value of its originality, that is, to *O*_*i*,ℭ_(*x*_0_), where *x*_0_ is the trait value at which the species *i* is more unique. Additionally, *Y*_*i*,ℭ_(*θ*) is homogeneous among species. In that sense, if the originality *O*_*i*,ℭ_ of species *i* is twice as large as the one of species *j* for all traits, *O*_*i*,ℭ_(*x*) = 2 *O*_*j*,ℭ_(*x*), the potential effect of species *i* is twice as large as the one of species *j* for all parameters *θ*: *Y*_*i*,ℭ_(*θ*) = 2 *Y*_*j*,ℭ_(*θ*). *Y*_*i*,ℭ_(*θ*) is also sublinear among species. This means that the potential effect of two species *i, j* considered together is less than or equal to (in cases where the two species functional space are identical) the sum of the potential effects of each one taken separately. For all *θ*,

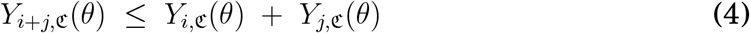

It is possible to understand the role of species *i* in the studied ecological process by examining its potential effect *Y*_*i*,ℭ_ variation in *θ*. The higher the overall value of the function, the greater the species effect on the process. When *θ* is equal (or close) to 1, *Y*_*i*,ℭ_(1), provides the average value of species *i* originality, *O*_*i*,ℭ_, weighted by the resources community functional space *C*_ℜ_. A large value of the potential effect indicates that species *i* participates in many interactions where the resources community functional space *C*_ℜ_ is large. In this case, the potential effect is more sensitive to modifications of the resources community. The picture for large values of *θ* is different. By increasing *θ*, a greater weight is given to interactions where the participation of the species *i* is major or even unique. Therefore, if the presence of the species is necessary or very important for the interaction to occur, the value of the species effect *Y*_*i*,ℭ_(*θ*) will be large. The potential effect of a species is not much sensitive to the resources community functional space *C*_ℜ_ for large values of *θ*.

A sharp increase of *Y*_*i*,ℭ_(*θ*), as *θ* varies from 1 to infinity, means that species *I* is highly original and that its originality reaches its maximum in a set of traits which are rare in the resources community. In contrast, the potential effect of species *i* remains stable with the increase of *θ* when it is not unique or when its originality reaches its maximum close to the region of traits where the resources community reaches its maximum.

### Niche amplitude

A species potential effect, *Y*_*i*,ℭ_(*θ*), is determined by many variables, including the niche amplitude, which plays an important role in the analysis of a species contribution to an ecological process. Here we introduce the species *i* niche amplitude, *W*_*i*_ as

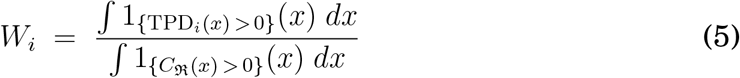

In this equation, 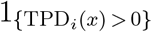 is the indicator function of the species *i* TPD_*s*_ and 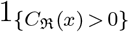 denotes the resource community functional space, *C*_ℜ_. For the consumer species *i*, the indicator function is equal to 1 when an interaction with trait value *x* is possible and equal to 0 when there is no interaction. For the resource community, the indicator function is equal to 1 when trait value *x* is present in the community and equal to 0 when it is not. Therefore, *W*_*i*_ stands for the range of trait values with which species *i* interacts with divided by the extent of the set of trait values available in the resource community. We divided by the range of trait values of the resource community so that the niche amplitude *W*_*i*_ is a number between 0 and 1.

By definition, generalist species have a large niche amplitude, and specialists a small one. If a generalist species has a small potential effect for *θ* = 1, it either has low originality or does not interact with frequent traits in the resources community. If it has high originality at some trait values, these resources availabilities are small. To investigate whether there are points of high originality one must compute the potential effect for high values of *θ*. A similar analysis can be achieved for a specialist species with a large potential effect for *θ* = 1. In this case, the species has high originality for traits that are common in the resources community. Therefore, for an accurate interpretation of the role of a species on an ecological process, both its potential effect *Y*_*i*,ℭ_(*θ*) and its niche amplitude *W*_*i*_have to be considered.

### 3.1 Examples

To exemplify our framework, in Figure 2 we detach 4 species with different trait distributions from a community composed by 10 species. We focus on the influence of species niche width and originality, and the resources community functional space. Therefore, in the examples we assume that all species’ niches have the same quantitative contribution to interactions, *V*_*i*,𝔞_*/V*_𝔞_ (see equation (2)), to exclude this variable influence on the species potential effects.

**Figure 2:**
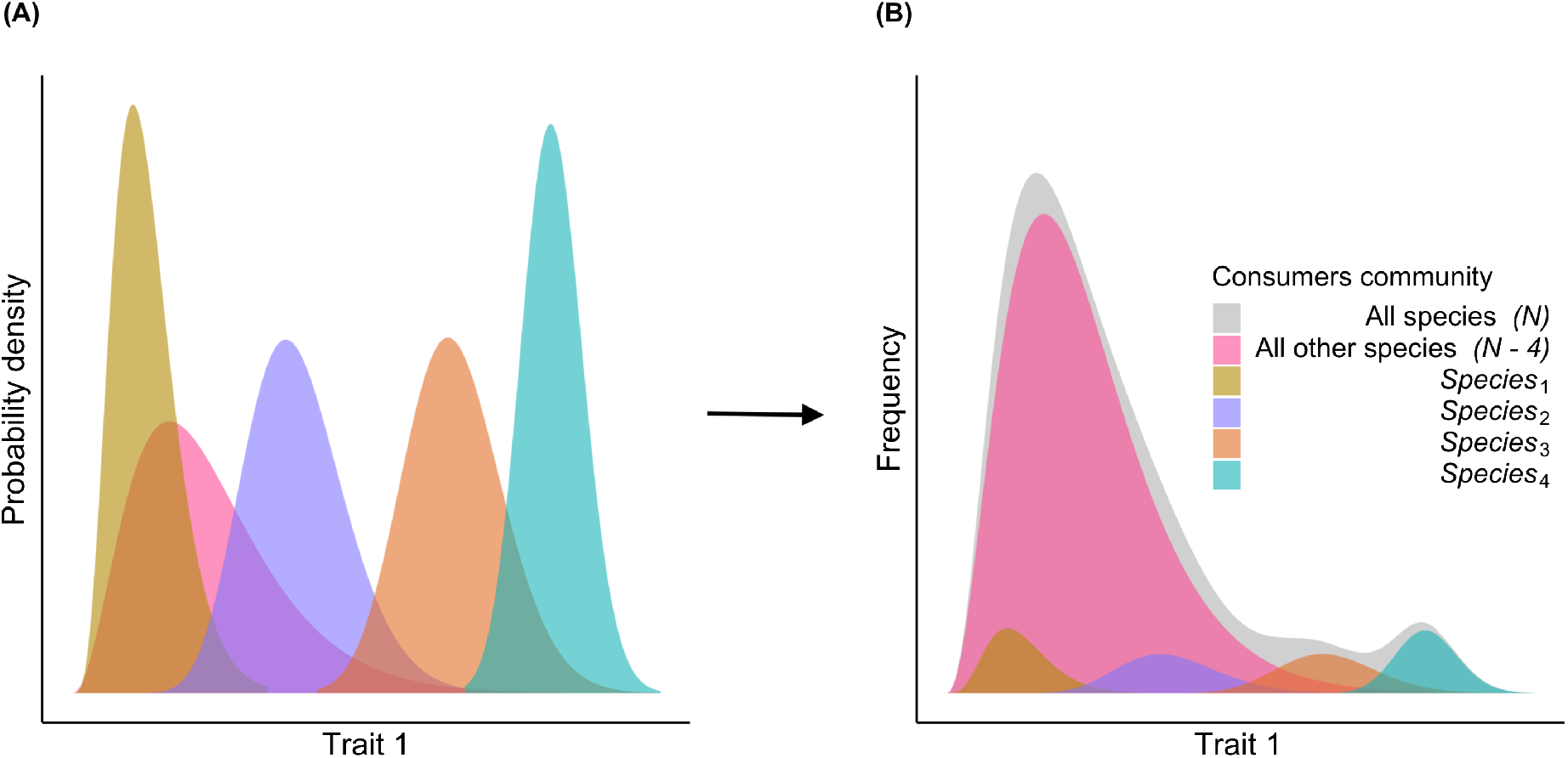
Consumers community functional space estimation. In **(A)** we detach 4 species TPD_s_ from the consumers community (N=10) to be used in the examples (see Figure 3 below). The TPD_s_ from the 6 other species that compose the community are represented in pink. In **(B)** the species TPD_s_ were balanced by their quantitative contributions to estimate their functional spaces. The species’ relative quantitative contribution can be represented by the number of interactions (see equation 1) or abundance (see equation 9). All species have a relative quantitative contribution of 0.1. The consumers community functional space is being represented in grey.

Each species has different ranges of trait values. Species 1 and 4 have a narrower niche, whereas species 2 and 3 niches are wider. Moreover, species 1 and 2 interact with trait values that are frequent within the rest of the consumer community, represented in pink, while species 3 and 4 interactions are rare or inexistent in the rest of the community.

In examples 1 and 2 (Figure 3), different resource availabilities are explored (Figure 3 **(A)** and **(C)**) to show the importance of including the resource community functional space into analysis. The scale adopted to plot the resource functional space is different from the consumers community’s and species’ functional space so it is possible to fit them in the same figure. Figure 3 **(B)** and **(D)** present species potential effects *Y*_*i*_ as a function of *θ*. As we mentioned above, *Y*_*i*_ increases with *θ*. Note that this doesn’t mean that species effect is related to *θ*. This parameter has been included in the model only to identify each species’ originality in interactions. On the left part of the plots, for *θ* close to 1, the value of the function *Y*_*i*_ gives species *i* potential effect without emphasizing its originality. In contrast, on the right, for large values of *θ*, the species effect is computed giving a significant weight to its originality.

**Figure 3:**
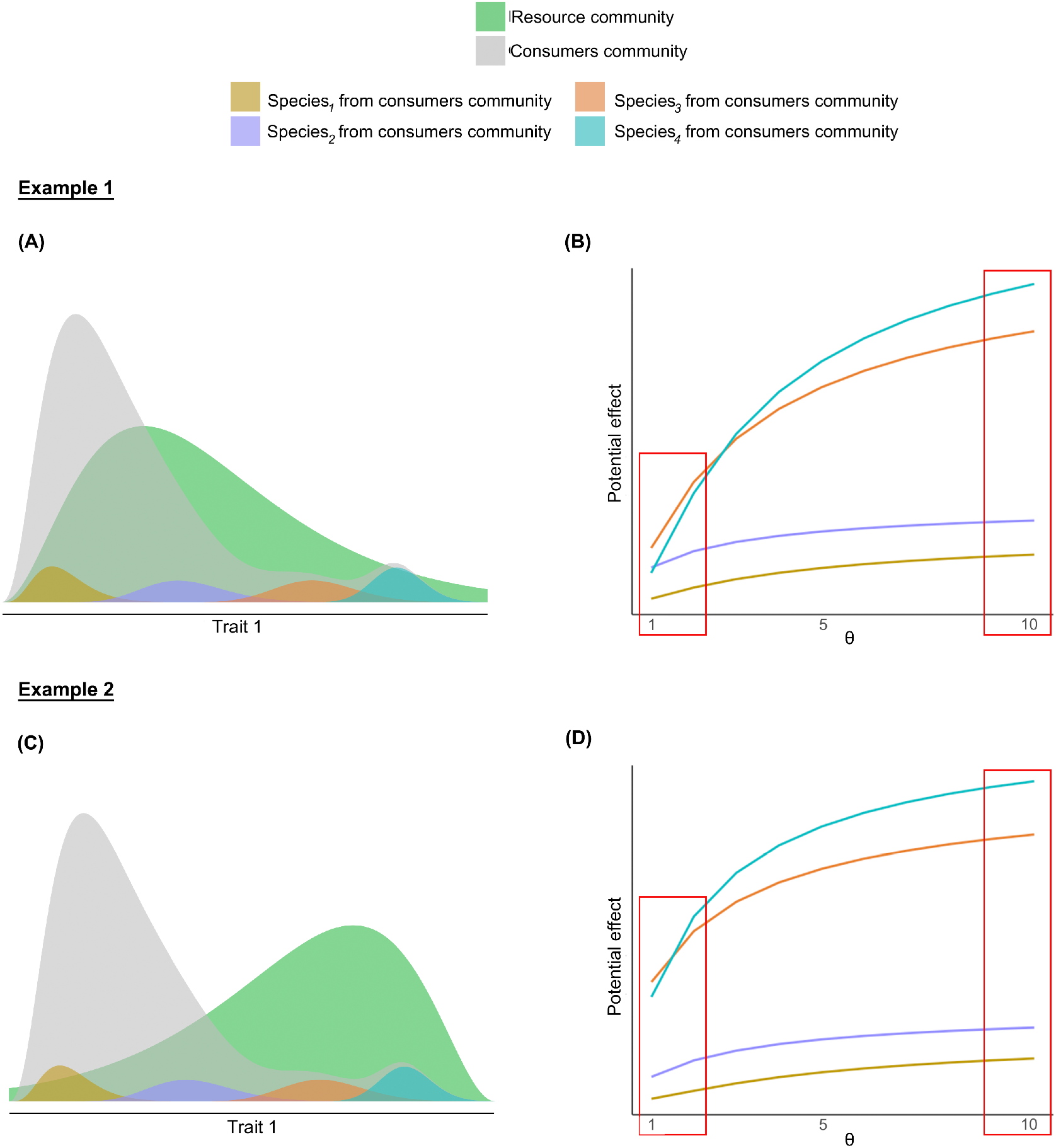
Species’ potential effects on trophic interactions. In example 1, the consumers and resources communities functional spaces have high densities for similar values of trait 1. In contrast, in example 2 the communities maximum densities are uneven. Species follow the same colour code depicted in Figure 2.

In the first example (Figure 3 **(A)**), species 1 is redundant and interacts with trait values that are not very common in the resource community *C*_ℜ_. Species 2 is less redundant and interacts with the most frequent traits on *C*_ℜ_. Species 3 is more unique than the other two and interacts with common resources trait values. Finally, species 4 is the most unique but interacts with rare trait values from *C*_ℜ_.

Species 2 and 3 have the highest potential effects on the ecological process *Y* when originality is given the same weight of the other variables, as indicated on the left portion of the plot in Figure 3 **(B)**. Species 3 has the highest impact because its originality takes large values on the set of trait values where *C*_ℜ_ also takes large values. Despite its redundancy, species 2 has the second highest value. That is because its niche extends through most of the region where the resources community has the highest probabilities. Species 1 and 4 potential effects are smaller because their niche range is concentrated in a region where the resources community density is small. The effect of species 4 is higher than species 1 because of its high originality. This analysis illustrates that in the evaluation of the potential effect *Y*_*i*_ of species *i*, both its originality and the agreement between its niche range and the density of the resources community *C*_ℜ_ are important.

On the right part of plot **(B)** (Figure 3), where *θ* is high and species originality is given a greater weight, species 4 has the highest potential effect. This is because species 4 is the most original species and is responsible for more unique interactions among the four species, as seen in plot **(A)** (Figure 2). Note that the potential effects of species 1 and 2, *Y*_1_ and *Y*_2_ respectively, almost do not vary with *θ*, meaning they are very redundant.

We included example 2 (Figure 3) to reinforce our claim that species with the same trait space and same relation to their communities functional space may have different effects on the ecological process as resource availability changes. We maintained the TPD_*s*_ of the consumer species from example 1 and reversed the resources community functional space. As the TPD_*s*_ of the consumer species were not modified, their originality did not change. However, in example 1 most species of the total consumers community, represented in grey, interacted with common trait values in the resource community. Meanwhile, in example 2 most frequent trait values in the resource community have a small probability of being consumed.

The effect of the modification of the resource community can be observed in Figure 3 **(D)**. Species 2 potential effect decreased for small values of *θ*, as in this case this species interacts mostly with rare trait values in the resources community. In contrast, species 4 potential effect increased for all values of *θ* as in this example it interacts with frequent trait values of the resource community. Observe that while the order of species potential effects were altered on the left part of the plot, it remains constant on the right. This happens because for large values of *θ* the species potential effect is more sensitive to modifications on originality than to changes in the resource’s functional space.

We also estimated each species’ niche amplitude. Species 2 and 3 have the highest amplitudes, *W*_2_ = *W*_3_ = 0.58, and species 1 and 4 the lowest ones, *W*_1_ = *W*_4_ = 0.33. Observe that species 3 and 4 have similar potential effects for *θ* = 1 although the former has a greater niche amplitude than the latter. This happens because species 4 is more original than species 3, as shown in the right of plot **(D)** (Figure 3). Likewise, species 2 and 3 have the same niche amplitude but species 3 potential effect is greater for *θ* = 1, as it interacts with more frequent trait values in the resource community and is more original than species 2.

These examples highlight the fact that the species’ effects on an ecological process is stronger when its originality is close to 1 and its functional space matches the one of the resource community. The wider the niche is, the greater the potential effect it might have on the process. The parameter *θ* allows to tune the importance of originality in the estimation of species potential effect.

### 3.2 Robustness

In this subsection, we examine the influence of species quantitative contribution to interactions on its potential effect. To illustrate this issue, we return to Example 1. In Figure 4 **(A)**, we estimate species 2 and 4 potential effects, *Y*_2_ and *Y*_4_, for different quantitative distributions. In Figure 4 **(B)** we compute species 4 potential effect removing species 3 from the consumers community, so that some trait values become strictly unique to species 4.

**Figure 4:**
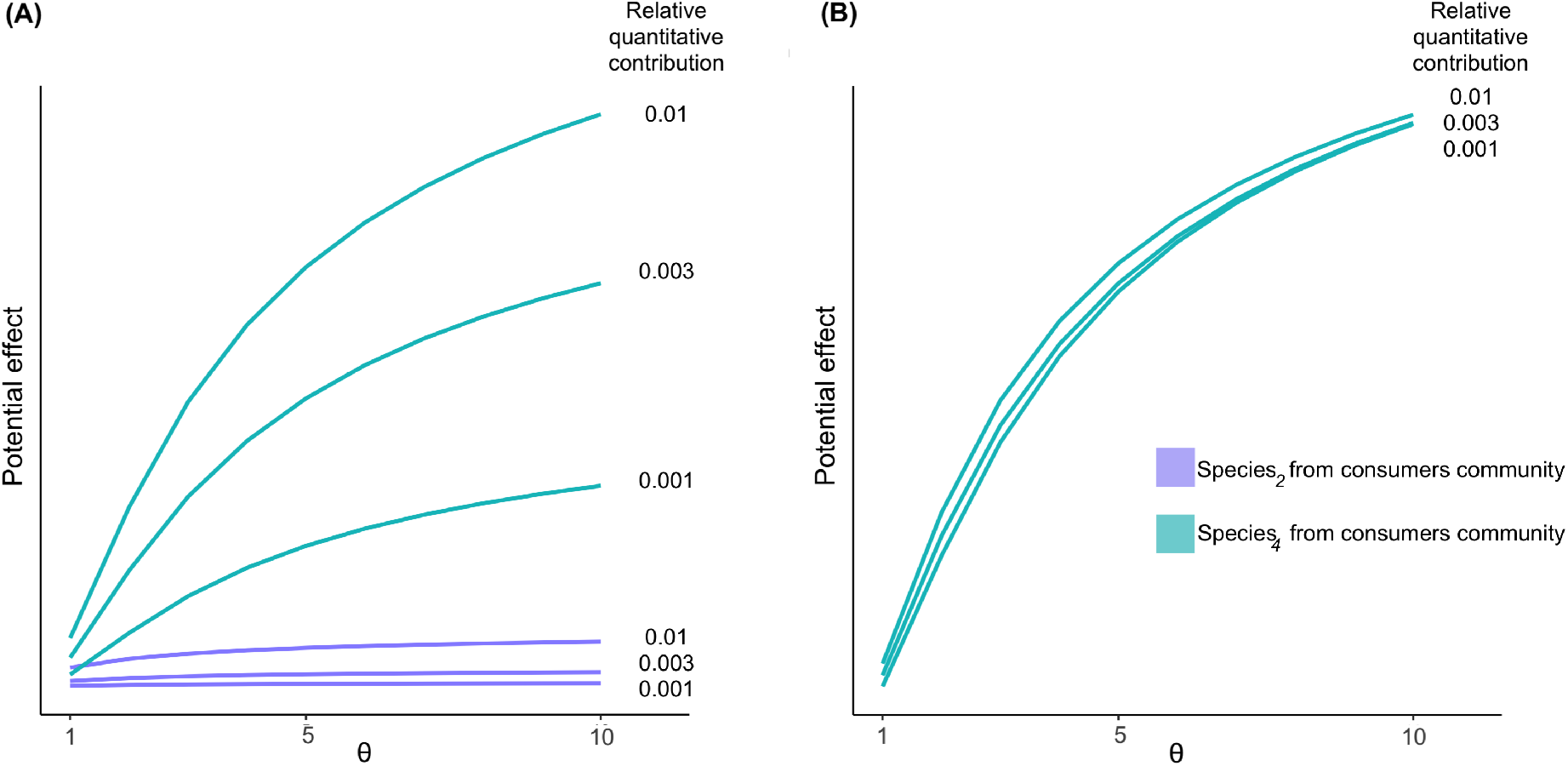
The influence of species relative quantitative contribution on its potential effect. In **(A)** species 2 and 4 depicted in Figure 3 **(A)** have their potential effects estimated for different relative abundances. Although the effect varies with the relative quantitative contribution, species 4’s potential effect is always higher than species 2’s. In **(B)** species 4 potential effects is estimated removing species 3 from the consumers community. In this case some of species 4’s trait values become strictly unique and the species’ potential effect almost does not vary with modifications in the relative volume contribution.

Figure 4 **(A)** shows that although the potential effect is sensitive to modifications of the quantitative variable, species 4 remains with a higher potential effect than species 2 in all cases. In contrast, Figure 4 **(B)** shows that when some trait values are strictly unique to the species, the potential effect is not very sensitive to changes in the species quantitative contribution. Therefore, predictions are more robust to variations in the quantitative variable when species have unique interactions. When species do not participate in unique interactions their effects change proportionally to modifications in their quantitative contribution and relations among species do not change.

Due to the framework’s sensitivity to quantitative data, researchers and practitioners must be cautious when interpreting the framework’s results. If the difference between species’ potential effects is small, it should be interesting to make tests with different relative quantitative contributions, or even remove the quantitative variable (see the Case Study section). Nevertheless, when species have unique interactions one can more freely interpret the results.

## 4 Framework modifications

In this section, we present modifications of the framework. Subsection 4.1 shows how to compare the potential effect between groups instead of species. Subsection 4.2 proposes to replace a species quantitative contribution to interactions introduced above in equations (1) and (2) by data easier to assess, such as species relative abundances and richness. Finally, subsection 4.3 provides two different ways to add an absent species in the versions present in subsection 4.2.

### 4.1 Groups potential effect

Occasionally, one might be interested in the potential effect of groups instead of species. In this case, the originality of a group *G*, denoted by *O*_*G*,ℭ_, is computed as the sum of its species originalities:

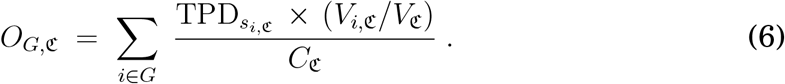

In this equation, the sum is calculated over all species *i* which belongs to the group *G*.

The communities functional space, given by equation (1), remains the same, while the potential effect of the group *G* is given as in equation (3), but with the species *i* originality *O*_*i*,ℭ_ replaced by the group’s *O*_*G*,ℭ_:

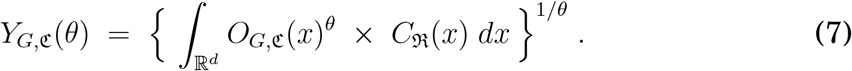

As already observed in equation (4), the potential effect of the group is smaller than the sum of the potential effects of the species that form the group:

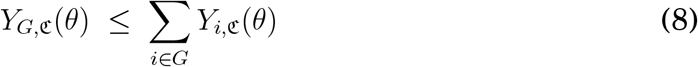

for all *θ* ≥ 1.

### 4.2 Replacing the quantitative contribution

A species contribution to their community’s functional space is defined by two elements: their trait probability density (TPD_*s*_) and their relative quantitative contribution (see equation (1)). In the conceptual model we define the quantitative variable as the relative contribution of species *i* to the total of interactions realized by its community. However, this kind of data is not available for many systems and processes derived from trophic interactions and can be replaced by proxies. If working with pollination, for instance, we could use standardized records of flower visits per species (as a proxy for pollination) so it would be possible to calculate the relative contribution of each species for the total number of flower visits performed by the pollinator community (Reynolds et al. 2009). We propose here two possibilities for when this data is not available. In the first one, the quantitative contribution is estimated by species relative abundance and in the second one it is assumed to be the same for all species. These approaches must be implemented cautiously because the potential effect is sensitive to modifications of the quantitative contribution, as discussed in Section 3.2.

#### Abundance as quantitative contribution

Abundance is an important factor determining the probability of species encounter and therefore, species interactions. In most network studies, the absence of interaction does not necessarily mean an impossible interaction, but that there were no records of such interaction (Bluthgen 2010). Abundant species often interact with many species and perform a large proportion of the realized interactions. In this way, it is reasonable to use species abundance as a proxy for species contribution to the total realized interactions (Canard et al. 2014).

In this modification, the species TPD_*s*_ is weighted by their relative abundance in their community *A*_*i*,𝔞_*/A*_𝔞_. Here, *A*_*i*,𝔞_ is the species abundance and *A*_𝔞_ the total abundance in the community. In this case, the community functional space *C*_𝔞_ is defined as:

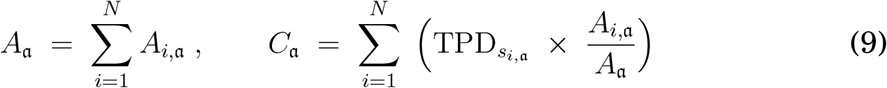

and the originality of species *i* represented by *O*_*i*,𝔞_ as:

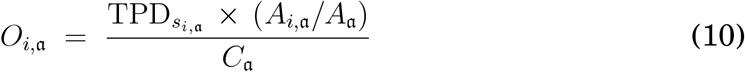

The potential effect of species *i* on the ecological process represented by *Y*_*i*,𝔞_(*θ*) is defined as showed in Equation (3).

#### Uniform quantitative contribution

Although abundance is important to estimate species’ effect on an ecosystem process (Suding et al. 2008), this information is not always available. Nevertheless, a species’ potential effect can still be estimated.

In many situations the only data available for researchers and practitioners is a list of species. In such cases, when the same quantitative contribution is assumed for all species, the species TPD_*s*_ is balanced by the total number of species *N* in the community (species richness). The functional space of the communities, denoted as *C*_𝔞_, is therefore defined by:

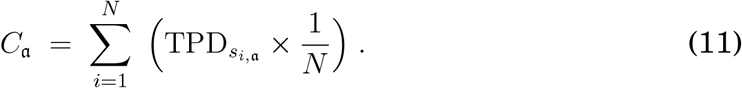

The originality of species *i* is given by

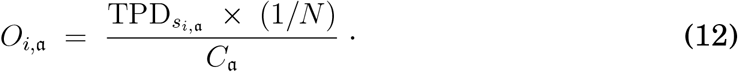

Note that this equation can be rewritten as

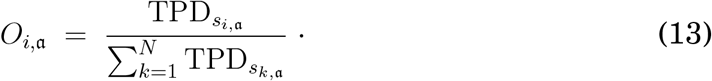

The potential effect *Y*_*i*,𝔞_(*θ*) of species *i* is given by equation (3).

One might argue that this model is oversimplified as species’ role in trophic interactions are often dependent on species abundance and density (Akçakaya et al. 2019). However, on the one hand, even more simplified models, which take into account only the convex hull of the niche, are commonly used (Cornwell et al. 2006) and effectively compares species roles on ecological processes (Quitián et al. 2019, Dehling et al. 2020). Additionally, traits affect both interactions and abundance (Bartomeus et al. 2016). Accordingly, species trait probability densities (TPD_*s*_) and similar approaches can give relevant informations about species effect on an ecological process (Kuppler et al. 2017, Dehling and Stouffer 2018, Cooke et al. 2019). For example, when species abundances are low, as it usually is the case in modified landscapes, species richness predicts well seed dispersal functionality (Rumeu et al. 2017).

### 4.3 Including absent species into the framework

In this subsection, we consider the case where one or more species are absent from the community. We assume that these species TPD_*s*_ can be estimated from the literature. We propose modifications in the models of the previous subsection to estimate the potential effect of these species if they were introduced to the community.

#### Abundance as quantitative contribution

Consider *N* the total number of species in the community and by *M* the number of absent species to be included in the model. Let *A*_*i*,𝔞_, 1 ≤ *i* ≤ *N*, be the abundance of the present species *i*. The abundance that the absent species *j* often show in similar environments is denoted as *A*_*j*,𝔪_, 1 ≤ *j* ≤ *M*. The functional space formed by the set of absent species, represented by *C*_𝔪_, is given by

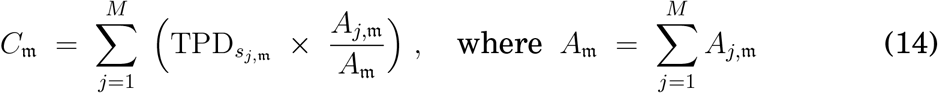

and 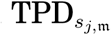 stands for the trait probability density of the absent species *j*. The total abundance and the functional space of the present species, represented by *A*_𝔞_ and *C*_𝔞_, respectively, are computed as in equation (9). The functional space of the total community, denoted by *C*_𝔞𝔪_ can be computed as a weighted average:

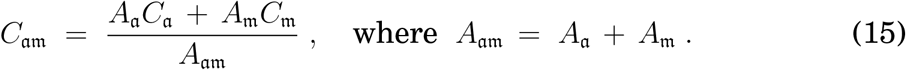

The originality of the absent species *j*, denoted by *O*_*j*,𝔪_, is given by

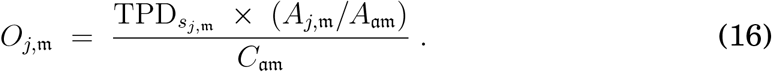

Once more, the species potential effect is estimated as in equation (3).

#### Uniform quantitative contribution

When there is no information about species abundance one can address the presence-based model from the previous subsection (equations (11) and (12)). In this case the functional space of the set of absent species *C*_𝔪_ and the total community functional space *C*_𝔞𝔪_ are determined as

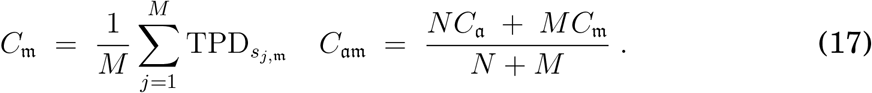

The absent species *j* originality is thus defined as

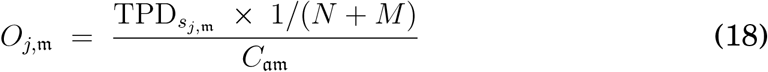

and the potential functional effect *Y*_*j*,𝔞𝔪_ as in equation (3).

## 5 Case study

We illustrate our framework using the seed dispersal network of Tijuca National Park (TNP, 3.953 ha), within the city of Rio de Janeiro. For centuries, most of the area that now corresponds to the Park had been exploited for coffe farming and coal production. In the mid 19^th^ century, it began to be reforested to restaurate the city’s water supply (Padua 2002). Because this park is surrounded by a metropolitan matrix, the missing fauna has been unable to recolonize TNP; many important original vertebrates are still absent from this area.

Refaunation (Oliveira-Santos and Fernandez 2010) and trophic rewilding (Svenning et al. 2016) have been introduced as solutions to reverse the effects of defaunation. They propose to restore ecosystems through the reintroduction of recently extirpated species. The main difference between approaches is that the latter prioritizes the reestablishment of interactions over the fauna per se. Nevertheless, the reestablishment of interactions has always been a central concern in the idea of refaunation. It was suggested as an alternative to pleistocene rewilding (Donlan et al. 2006) (do not confound pleistocene rewilding with trophic rewilding), so only recently extinct native species would be reintroduced and there would be a lower chance to provoke unwanted interactions. Here, they will be considered as equivalent terms and refaunation will be used when we refer to these propositions.

In 2010, a refaunation project started in the TNP, aiming to reintroduce missing species and reestablish ecological processes affected by their local extinction (Fernandez et al. 2017). Since then, agoutis (*Dasyprocta leporina*) and howler monkeys (*Alouatta guariba*) have been reintroduced. However, to date, there are no studies comparing their contribution to seed dispersal considering the present frugivores community and resource availability.

In this case study, we 1) analyse the role in seed dispersal of the groups of frugivores present in the Park before the refaunation and 2) compare the potential effects of the reintroduced species. In the future, it would be interesting to use this framework to assist in the selection of new species to be reintroduced and the order they should be reintroduced in.

For this purpose, we used data from the Atlantic Frugivory data set (Bello et al. 2017) to build the TNP seed dispersal network. Next, we constructed the frugivore and plant species TPD_*s*_, as described in the Methods section, based on fruit and seed length and diameter. Then, we removed seed length, as a PCA showed a strong correlation among variables (*R* = 0.9).

As we did not have information about the frequency of interactions or the abundance of species, we adopted the presence-based model (equations (11) and (12)) to obtain the frugivores’ functional spaces and the originality of species. We used equation (6) to compute the originality of each frugivore group (with *V*_*i*,ℭ_ = 1 for all species *i*). To compare the potential effects of the reintroduced species, we considered two different scenarios. In the first one, we added agoutis to the community and in the second one we added howlers. In both, we adopted equations (17) and (18) with *M* = 1. All potential effects were estimated through equation (3).

Figure 5 **(A)** presents TNP’s frugivores potential effects grouped into taxonomical orders and Figure 5 **(B)** compares the potential effect of agoutis and howler-monkeys in TNP’s seed dispersal. To study robustness in practical examples, in Figure 5 **(C)** we gave each frugivore group weight equal to 1, and in Figure 5 **(D)** we decreased the agoutis’ weight to 0.1, but kept howlers weight equal to 1.

**Figure 5:**
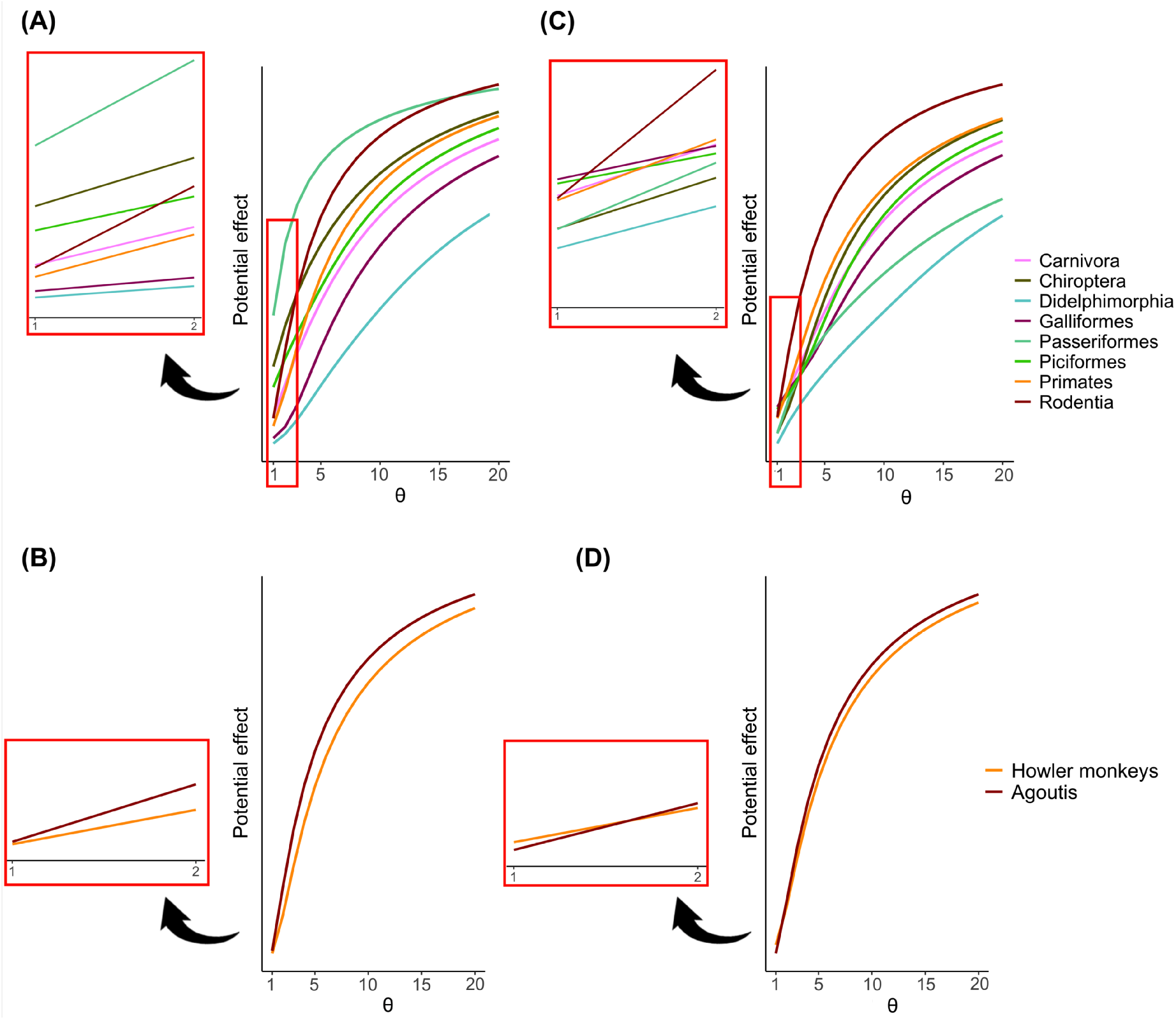
Groups and reintroduced species potential effects in Tijuca National Park seed dispersal network. **(A)** and **(C)** represent the remaining frugivores separated by taxonomic groups. In **(A)**, a group’s quantitative contribution to the interactions is given by the number of species in their composition. In **(C)** we gave all groups the same weight to comprehend the disadvantages of using the presence-based model. **(B)** and **(D)** depicts the comparison between the reintroduced frugivores’ potential effect. In **(B)** agoutis and howler monkeys have the same weight. In **(D)** we gave agoutis weight 0.1, while howlers remained with weight 1, so we could explore how the species potential effects would change if they had different relative abundances.

The analysis for the frugivores groups (Figure 5 **(A)**) demonstrates that Passeriformes are dominant for most values of *θ*, while Galliformes and Didelphimorphia have the lowest potential effects for any value of *θ*. Indeed, while Passeriformes is composed by 86 species, Didelphimorphia and Galliformes are represented by only one species. In contrast, although Rodentia is also represented by one species, it has the highest potential effect for large values of *θ* due to its high originality. For the same reason, Primates’ potential effect also rises significantly with the increase of *θ*. Note that the potential effects of the different orders are substantially different for *θ* close to 1 and for high values of *θ*. Therefore, the potential effect analysis of a species or group can differ if one is specially interested or not on groups originality.

Figure 5 **(B)** shows that when agoutis and howlers have the same volume contribution, agoutis have a higher potential effect for all values of *θ*. However, if one does not value unique interactions, the species effects are similar. Agoutis also presented a greater niche amplitude (*W*_*agouti*_ = 0.99 and *W*_*howler*_ = 0.27), meaning that agoutis are more generalists than howlers. Since agoutis have a greater niche amplitude than howlers, but similar potential effect for *θ* = 1, the howlers’ TPDs is greater than the agoutis’ for frequent trait values on the resource community. Therefore, while howlers interact more often with frequent fruit trait values in the plant community, agoutis add rare interactions to the TNP seed dispersal network and interact with a broader set of resource trait values.

In Figure 5 **(C)** and **(D)** we altered the groups and species quantitative contributions to illustrate the framework’s sensitivity to modifications in the quantitative variable, shown in Subsection 3.2. Some groups’ potential effect changed when the same weight was given to all groups (Figure 5 **(C)**). The greatest changes were observed in the groups composed by a higher number of species: Passeriformes, composed by 86 species, and Chiroptera, 18. The potential effect of Passeriformes was substantially reduced, as it became the second group with the smallest potential effect for all values of *θ*. Therefore, Passeriformes’s high potential effect in Figure 5 **(A)** is mainly due to the group’s high quantitative contribution. In contrast, the effects of Chiroptera were different for different values of *θ*. For *θ* close to 1, the group’s potential effect was markedly reduced. For high values of *θ*, its position almost did not change in relation to the other groups. Thus, Chiroptera is very original in both situations. Comparing plots **(A)** and **(C)**, it is possible to see that the relation between the groups potential effects was less modified in the right side of the plots than in the left. In the case of the reintroduced species (Figure 5 **(D)**), when we decreased the contribution of agoutis the order of the potential effect of the two species changed for low values of *θ*. Nonetheless, the species contribution remained the same for *θ* large. These results confirm the framework’s susceptibility to the quantitative variable for small values of *θ* and its robustness for large values of *θ*. It is also confirms that the model is more robust for more original species, due to the different variations of Passeriformes and Chiroptera.

The seed dispersal network from TNP has been intensely modified since its deforestation. Most of its large seed-dispersers, such as the muriqui (*Brachyteles arachnoides*) and the lowland tapir (*Tapirus terrestris*), were extirpated (Macedo 2017). The absence of these species has had an impact on large seeded plants that depend on larger animals to be dispersed. Moreover, some of these animals would not be able to be reintroduced due to their habits. The lowland tapir, for example, has a predominantly frugivorous diet and requires forest fragments larger than TNP’s area (Beca et al. 2017).

In this context, species need to be carefully chosen to efficiently reestablish lost functions. Although the framework here proposed informs on the roles of groups and species of seed dispersers, the variables used to determine their potential effects are not the only factors defining their functions. Muriquis and lowland tapirs, for example, are both dispersers of large seeds but diverge in relation to their seed deposition pattern (Bueno et al. 2013), which has an effect on post-dispersal seed fate and seedling recruitment (Lugon et al. 2017). Thus, we suggest that qualitative information on the consumers’ roles should be used to complement results on their potential effects.

Agoutis are good candidates to reestablish large frugivores roles. Our results show they are highly original and disperse larger seeds that have fewer dispersers in the Park. A recent study has identified that one of the largest seeds in TNP (*Joanesia princeps*) only germinated when buried by agoutis (Mittelman et al. 2020). Moreover, due to their scatter-hoader behaviour, agoutis are able to disperse large seeds over great distances (Jansen et al. 2012). Howler monkeys, on the other hand, do not disperse the largest seeds present in the Park. However, they interact with frequent traits in the resource community and therefore have an important role in TNP’s seed dispersal. Additionally, coprophagous beetles (Scarabeidae: Scarabaeinae) are associated with howlers feces (Genes et al. 2018) and can accidentally relocate the seeds present, increasing seed survival (Shepherd and Chapman 1998).

## 6 Discussion

As we have shown in the case study of seed dispersal, our framework can be applied using only species’ presence data. The results give powerful insights on the roles of each group participating in this particular ecological process, as well as important information on the potential effect of reintroduced species. The variation of the potential effect (*Y*_*i*,ℭ_) when varying *θ* combined with the niche amplitude (*W*_*i*_) informs on the type of the contribution: if it is due to the species or group quantitative contribution, originality of interactions or frequency of trait values with which they interact. For instance, the conclusion of reintroduced species’ analysis was that agoutis interact with a broader set of resource trait values and add rare interactions, while howlers interact more often with frequent resource trait values. Agoutis interact with a broader set of resource trait values because their niche amplitude is greater than howlers’; they add rare interactions because their potential effect is greater than howlers’ for large values of *θ* when the two species are given the same quantitative contribution; and, finally, howlers interact more often with frequent resource trait values because, despite the agoutis’ characteristics mentioned before, howlers’ and agoutis’ potential effects are very similar for *θ* close to 1.

Refaunation projects with the goal to mitigate the effects of species extinction are recently increasing (Seddon and Armstrong 2016). However, to date, there are no studies predicting species effect on ecological processes in refaunated areas. This kind of prediction, as given by our framework, is crucial for managers for choosing species for reintroduction, allowing them to decide between adding missing interactions or increasing resilience by adding redundant interactions. Reintroductions are very costly and time consuming, therefore, knowing beforehand the potential effects of species on ecological processes can help to choose which species to reintroduce and in which sequence. Additionally, our framework can also complement SWOT (strengths, weaknesses, opportunities, threats) analysis for reintroductions (White et al. 2015) and indicate areas that would benefit the most with certain species, by simulating the effect of reintroductions in different consumers and resource communities.

Genes et al. (2017) proposed a conceptual framework to evaluate the success of reintroductions regarding the reestablishment of ecological processes. The authors considered that just as species extinctions leave a debt of ecological interactions, their reintroduction has a credit of possible interactions to be restored. The success of the reintroduction would thus be reached when the reintroduced species re-established a pre-defined proportion or the totality of the amount of interactions it was expected to realize. This approach is entirely based on the quantity of interactions, represented by the number of species a reintroduced animal is expected to interact with. Our functional framework advances on this approach by adding a qualitative aspect to this assessment, as it informs which part of the resource trait space a reintroduced species is expected to fill (or interact with). This also allows to infer if the reintroduced species will interact with resources which are not used by the local consumer community, or, on the contrary, if the reintroduced species is expected to feed on resources that are already used by the local consumer community.

Studying species effects on ecological processes helps to understand the consequences of changes in community composition to the ecosystem functioning. Our framework can be applied with the opposite perspective of reintroductions to evaluate the possible effects of future extinctions and the consequences of past extinctions to ecological processes. The effect of extinctions is not random, as species differ in their response to different environmental drivers (Larsen et al. 2005). Likewise, species also differ in their effect on ecosystem processes (Suding et al. 2008) and consequences of extinctions on ecological processes will depend on the complementarity of species roles (Fründ et al. 2013). This means that the extinction of more original species is expected to result in stronger effects on ecosystem functioning. Nevertheless, redundancy is important for stabilizing communities (Pillar et al. 2013, Peralta et al. 2014) and prevents secondary extinctions in food webs (Sanders et al. 2018).

Our framework can also be used to analyse the impact of invasive exotic species on ecological processes. As suggested by Finerty et al. (2016), for predicting the effect of exotic species on ecosystem functioning we need to know if this species is able to shift the community trait space. The effects of biological invasion are not necessarily negative. In New Zealand, European exotic birds contributed to increase the generalism and consequently the effectiveness of seed-dispersal (Garcia et al. 2014). Therefore, this framework helps to examine if exotic species are contributing for a given ecological process or if they can be a risk for native species.

Despite the fact that species participate in different ecological and ecosystem processes (Hector and Bagchi 2007), our framework only allows for the analysis of one focal ecological process at a time. A species or group of species may have a weak potential effect ona given ecological process but strongly determine other processes. For example, invasive feral pigs (*Sus scrofa*) restored to some extent the dispersal of large fruits and seeds that were lost with the megafaunal extinction in the Pantanal (Donatti et al. 2011). However, due to their rooting behavior, they can reduce plant biomass, altering plant species composition (Barrios-Garcia et al. 2014). Therefore, evaluating species based on their potential effect on a single ecological process or service can undermine species importance (Hiron et al. 2018). When using this framework, it is essential to keep in mind that only one process is being accessed at a time.

Our framework does not account for species phenology. In fact, phenology is rarely considered in interaction networks studies, although it explains to a great extent properties of plant-animal mutualistic trophic interactions (Encinas-Viso et al. 2012). Nonetheless, the significance of species phenology in determining species interaction depends on the type of interaction. For instance, while phenological overlap is an important factor determining pollination (Peralta et al. 2020), for seed dispersal, interaction probability is better explained by trait-matching (González-Castro et al. 2015, Dehling et al. 2016). In the future, it would be interesting to incorporate species phenology into frameworks.

When both response and effect traits are considered, it is possible to predict how the community will change and its consequences to the ecosystem functioning. However, a multi-taxa perspective makes it difficult to aggregate response traits into the same analysis, as different taxa can respond through a wide variety of traits to environmental and anthropogenic drivers (Nowak et al. 2019). Nevertheless, diet can be considered a key functional trait, as it can both respond to external drivers and affect ecological processes (Hevia et al. 2017). For example, fragmentation often results in a frugivore community with smaller body size, which disperse smaller seeds in shorter distances (Schleuning et al. 2015). In turn, shifts in frugivores community functional composition can result in changes in trait values of the resource community (Galetti et al. 2013), with consequences in other ecosystem processes. In tropical forests, the selection of plants with small seeds by the extinction of large seed dispersers indicates a possible reduction of carbon storage capacity, due to the relation between seed diameter and carbon storage-related traits (Bello et al. 2015).

In a changing world, understanding the effects of changes in community composition is fundamental. In this context, our framework comprise an useful tool to predict how species extinctions, invasions and reintroductions will affect ecological processes resulting from trophic interactions. The framework synthesizes quantitative and qualitative information about the role of species in ecological processes. We believe the models presented here can be used as decision-making tools to improve invasive species management, species selection to restore ecological interactions and, consequently, enhance the ability to restore ecosystems.

